# Auditory biases in cognitive assessment – Insights from hearing-loss simulation for dementia screening

**DOI:** 10.1101/2022.10.05.510931

**Authors:** Christian Füllgrabe

## Abstract

Cognitive-screening tests are used to detect pathological changes in mental abilities. Many use orally presented instructions and test items. Hence, hearing loss (HL), whose prevalence increases with age, may bias cognitive-test performance in the target population for dementia screening. To study the effect of the test format, an impairment-simulation approach was used in normal-hearing listeners to compare performance on the Hopkins Verbal Learning Test when test items were unprocessed and processed to simulate age-related HL. Immediate verbal recall declined with simulated HL, suggesting that auditory factors are confounding variables in cognitive assessment and result in the underestimation of cognitive functioning.

## INTRODUCTION

To screen for dementia, verbal-learning tasks are used to detect changes in memory function [1]. One such screening test is the Hopkins Verbal Learning Test (HVLT) [2], in which a list of words must be remembered for recall. In addition to having good psychometric properties and diagnostic accuracy [3], the HVLT is easy to administer as instructions and test items are presented orally [2]. However, people screened for dementia are mainly aged over 65 years [4], with one third of this population being affected by disabling hearing loss (HL) [5], and nearly everyone in this age group showing some degree of decline in hearing sensitivity relative to young adults [6].

It has long been shown that persons with a HL perform worse on a variety of auditorily administered cognitive assessments than their normal-hearing (NH) counterparts [7, 8]. However, the exact reason for this observation is unclear. Poorer test outcomes could reflect the consequences of neuroplastic changes in the brain in response to prolonged impoverished auditory input [9, 10]. On the other hand, lower-than-normal cognitive performance could result from perceptual difficulties with the test format (i.e., reduced intelligibility of the auditorily presented instructions and test items and/or increased cognitive effort to process these auditory signals) [11, 12]. Both factors impact cognitive-test performance: while the latter has temporary effects during the assessment, and, thus, could be avoided by adapting the test format to the perceptual needs of the test person [13, 14], the former has more permanent effects on the neural substrate underpinning cognition which might only be amenable to long-term auditory rehabilitation (such as that provided by hearing aids) [15, 16]. When cognitively assessing older persons with HL using auditory screening tests (such as the HVLT) both factors may coexist and cannot be disentangled. To isolate the effect of HL during test administration, auditory deficits must be simulated in NH persons with intact cognition [17]. If a reduction in cognitive performance is observed in this test condition for these participants, the impact of the presentation format of the test is demonstrated. In the present study, such an impairment-simulation approach was used to test whether HL affects performance on the HVLT in terms of immediate verbal recall.

## MATERIALS AND METHODS

Thirty (10 females, 20 males) young (aged 18-23 years) native-English-speaking undergraduate students from Loughborough University (UK) with NH (defined as audiometric thresholds ≤ 20 dB Hearing Level at octave frequencies between 0.25 and 8 kHz) in the test ear were each assessed in three listening conditions. Lists 1-3 of the HVLT (each composed of 12 words) were recorded digitally (using a 44.1-kHz sampling rate and 32-bit quantization) from a female native-British speaker with a standard accent. The recordings were either left unprocessed to represent NH, or processed through a HL simulator [18] implemented in Matlab^®^ to mimic the following perceptual consequences of age-related HL (ARHL): loss of audibility (by attenuating the frequency components in several frequency bands), reduced frequency selectivity (by spectrally smearing the speech signal [19]), and loudness recruitment (by expanding the range of the speech signal’s envelope [20]). Two levels of HL severity were simulated based on epidemiological data [21]: a “mild” and a “moderate” ARHL, as experienced by the average 70- and 85-year-old, respectively [22]. The corresponding audiograms (see Figure 1A) were used as the input to the HL simulator.

**Figure 1.**
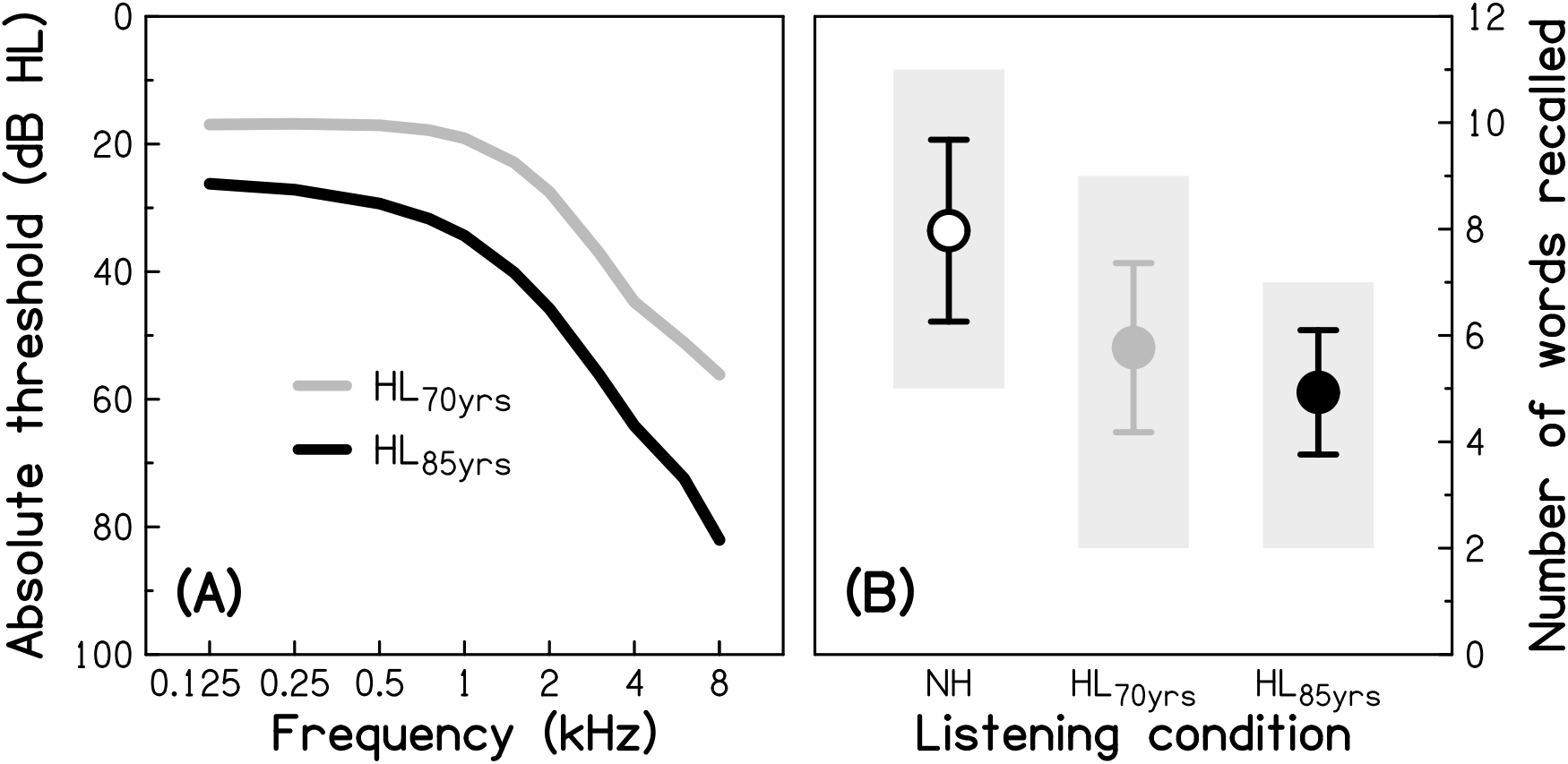
Audiograms representing the different levels of simulated age-related hearing loss (Panel A) and immediate verbal recall scores for each listening condition (Panel B). Absolute thresholds (in dB Hearing Level, HL) for the average 70-year-old (HL_70yrs_) and 85-year-old (HL_85yrs_), as used for the input to the hearing-loss simulator. Mean performance in the three listening conditions, using unprocessed stimuli to simulate normal hearing (NH) and processed stimuli to simulate age-related hearing loss experienced at two older ages (HL_70yrs_ and HL_85yrs_). Error bars represent ±1 standard deviation. Grey-shaded areas indicate the range of scores.

Auditory stimuli were presented over the right earpiece of Sennheiser HD 25 headphones to the participant seated in a quiet room. For the unprocessed stimuli, the presentation level was 70 dB Sound Pressure Level (corresponding to a raised conversational level presumably used when speaking to an older patient). Processed stimuli were presented using the same volume settings. Immediate recall after a single presentation of a given 12-word list was assessed. All analyses were performed using IBM SPSS Statistics Version 24 (IBM Corp., Armonk, NY). The study was reviewed and approved by Loughborough University Ethics Approvals (Human Participants) Sub-Committee. All participants provided written informed consent prior to taking part in the study.

## RESULTS

For the audiometrically NH participants in the unprocessed condition, mean performance in terms of the number of words recalled (*m* = 7.9) was similar to that previously reported for a similarly sized sample of young (but audiometrically unscreened) undergraduate students (*m* = 7.1) [23]. In the present study, mean performance (see Figure 1B) varied significantly across the three listening conditions (ANOVA: *F*[1.54, 44.6] = 42.74; *p* < 0.05, η^2^ = 0.596), and was significantly lower in each of the two simulated-HL conditions (for HL_70yrs_: *m* = 5.7, *p* < 0.001, 2-sided, Cohen’s *d* = 0.97; for HL_85yrs_: *m* = 4.9; *p* < 0.001, 2-sided, Cohen’s *d* = 1.66) compared to the NH condition. There was also a significant decline with increasing severity of simulated HL (*p* = 0.007, 2-sided, Cohen’s *d* = 0.61). The differences between conditions remained significant after applying a Bonferroni correction for multiple comparisons. For the most severe level of HL simulated here (i.e., a moderate HL), the number of words recalled dropped by three words (a reduction in recall performance by 38%). These results are consistent with findings from previous investigations of the impact of simulated HL on the performance on different cognitive tasks, albeit implementing cruder impairment simulations, such as physical attenuation (by using earmuffs) [24] or digital attenuation of the audio signal [25].

## DISCUSSION

The aim of the present study was to provide proof of concept that the perceptual consequences of ARHL interact with the presentation format of the HVLT, an orally administered verbal-learning task used to screen for dementia. To exclude any influences of neurological changes in response to long-term sensory deprivation on test performance, ARHL was simulated in young participants free of auditory and cognitive deficits.

A large, significant decline in performance with increasing simulated HL was observed. Since the same participants were tested in all listening conditions (following a within-subject design), this finding cannot be imputed to differences in cognitive functioning, but reflects, at least partially, the consequence of reduced audibility: test items that are not (or only partially) heard cannot be recalled (correctly). In clinical practice, “adequate” audibility could be verified after test administration by asking the patient to verbally identify each test item. However, being able to recognize the test items does not mean that cognitive performance is not affected [17, 26]. Indeed, age- or HL-related changes in suprathreshold auditory processing abilities (that are audibility unrelated) lead to distortions in the internal representation of the auditory signal. It is assumed that this requires the listener to use cognitive resources to achieve speech recognition [11, 27]. Consequently, fewer cognitive resources are available for the completion of the cognitive task itself, resulting in lower test performance. Importantly, this means that measures implemented by clinicians to ensure “fair” test conditions for older patients (such as speaking louder during test administration or letting the patient wear hearing aids) at best improve audibility but do not compensate for increased listening effort associated with perceptual distortions.

In addition to affecting performance scores, (simulated) HL may also change the composition of the cognitive processes associated with the completion of a cognitive task [26]. Hence, the comparison of test outcomes from NH participants and participants with HL and the interpretation of observed differences in terms of a decline in specific cognitive processes might be flawed.

Several caveats regarding the reported findings should be noted: First, a relatively small number of participants was tested in the present study. However, the observation of a significant effect of simulated HL on cognitive-test performance echoes results from previous impairment-simulation studies investigating other cognitive tests using smaller and larger sample sizes [17, 24, 26, 28-30]. Second, only the effect of simulated HL on immediate verbal recall after a single presentation of the word list was assessed; delayed recall and recognition, which are also part of the revised version of the HLVT [31] and often used as independent or complementary indicators for changes in memory functions, were not explored. Finally, the HL simulator mimicked only some of the perceptual consequences of ARHL; for example, the decline in temporal processing abilities that also occurs with ageing and HL [32, 33], and that has been shown to be associated with speech intelligibility [34, 35], was not simulated. Hence, it is likely that the present study underestimated the extent of the impact of ARHL on verbal-memory performance and that the true auditory bias in cognitive assessment is even larger than that reported here.

In summary, the results of the present impairment-simulation study suggest, by extrapolation, that in older patients who are screened for dementia using verbal-learning tasks (such as the HVLT), hearing status may be a confounding variable. As the prevalence of ARHL is high in the target population for dementia screening, there may be a considerable risk of mis- or over-diagnosing cognitive decline when using auditorily presented cognitive tasks [17, 36].

To avoid such biases, several of the most frequently administered cognitive screening tests have been adapted in recent years to better fit the needs of people with HL [37-39]. In most cases, this simply meant converting the auditorily presented instructions and test items of the standard version into visual material. However, the uptake of these modified versions is relatively low, and a formal validation is generally lacking [40]. In addition, evidence that the visual form of a given test yields less biased and thus higher performance in people with HL is mixed [25, 37, 41]. It is conceivable that by presenting the cognitive task in the visual domain, the bias issue is simply shifted to another sensory modality, as visual processing is also affected by the aging process [42, 43]. Thus, it currently seems premature to make firm recommendations how best to assess cognitive abilities in people with HL.

In conclusion, it is hoped this study will help raising awareness amongst clinicians and researchers of the existence of sensory biases in cognitive assessment [17, 36] and may contribute to improving clinical screening and assessment tools [44].

## ACKNOWLEDGEMENTS

The author is grateful to Magdalena Margol-Gromad for help with recording the test stimuli, Lionel Fontan for support with the hearing-loss simulation, and Bleron Mjekiqi for help with data collection. The author also thanks Tom Baer for his insightful discussions, and three anonymous reviewers for constructive feedback on earlier versions of the manuscript.

## CONFLICT OF INTEREST

The author has no conflict of interest to report.

## Notes

### Competing Interest Statement

The authors have declared no competing interest.

